# Awake mouse brain photoacoustic and optical imaging through a transparent ultrasound cranial window

**DOI:** 10.1101/2021.12.08.471795

**Authors:** Shubham Mirg, Haoyang Chen, Kevin L. Turner, Jinyun Liu, Bruce J. Gluckman, Patrick J. Drew, Sri-Rajasekhar Kothapalli

**Affiliations:** Department of Biomedical Engineering, Pennsylvania State University, University Park, PA 16802, USA; Penn State Cancer Institute, Pennsylvania State University, Hershey, PA 17033, USA; Graduate Program in Acoustics, Pennsylvania State University, University Park, PA 16802, USA; Department of Engineering Science and Mechanics, Pennsylvania State University, University Park, PA 16802, USA; Department of Neurosurgery, Pennsylvania State University, University Park, PA 16802, USA; Center for Neural Engineering, Pennsylvania State University, University Park, PA 16802, USA

## Abstract

Optical resolution photoacoustic microscopy (OR-PAM) can map the cerebral vasculature at capillary level resolution. However, the OR-PAM setup’s bulky imaging head makes awake mouse brain imaging challenging and inhibits its integration with other optical neuroimaging modalities. Moreover, the glass cranial windows used for optical microscopy are unsuitable for OR-PAM due to the acoustic impedance mismatch between the glass plate and the tissue. To overcome these challenges, we propose a lithium niobate based transparent ultrasound transducer (TUT) as a cranial window on a thinned mouse skull. The TUT cranial window simplifies the imaging head considerably due to its dual functionality as an optical window and ultrasound transducer. The window remains stable for six weeks, with no noticeable inflammation and minimal bone regrowth. The TUT window’s potential is demonstrated by imaging the awake mouse cerebral vasculature using OR-PAM, intrinsic optical signal imaging and two-photon microscopy. The TUT cranial window can potentially also be used for ultrasound stimulation and simultaneous multimodal imaging of the awake mouse brain.

---

Optical resolution photoacoustic microscopy (OR-PAM) is a capable technique for quantitatively assessing cerebral microvasculature and holds significant potential for studying various patho-physiological processes in the brain [1]. It is based on the photoacoustic principle where photon absorption leads to emanation of broadband acoustic waves in chromophores such as melanin, lipids, oxy- and de-oxy hemoglobin [1]. OR-PAM has been extensively used to study vasculature as it can provide multiparametric information including hemoglobin concentration, blood flow, blood oxygenation as well as the metabolic rate of oxygen [2, 3]. Moreover, OR-PAM can also leverage neurovascular coupling to assess neural activity as a correlate of hemodynamic responses [4]. Its ability to map microvasculature with sub-micron to micron resolution makes it a suitable technique for studying dysfunction in neuro-microvasculature linked to various brain physiological and pathological processes such as aging, Alzheimer’s, multiple sclerosis, tumor growth and other neurological disorders [1].

Previous demonstrations of OR-PAM relied on imaging rodent brain in anaesthetized states [5]. However, as anesthesia induces noticeable changes in the brain [6], the field has shifted towards monitoring hemodynamic changes in awake animals. Consequently, this has brought various engineering challenges to accommodate the rodent’s awake state using either head mounted or head restraint setups. Head mounted setups [4] for OR-PAM are able to image mouse brain in freely moving conditions, but are limited by a small field of view as well as long term mechanical and optical stability considerations. Head restraint setups, on the other hand, allow for a larger field of view and exhibit long-term stability [2, 3]. Another challenge for OR-PAM brain imaging setups is to overcome scattering by the skull which limits imaging resolution and penetration depth. Conventionally, OR-PAM brain imaging setups have relied on craniotomies [4] or skull thinning [2, 3] to overcome or reduce scattering. However, craniotomy can potentially induce inflammatory response leading to alterations in neural processes and blood flow [7]. On the other hand, skull thinning has minimal inflammation but suffers from bone regrowth [7]. Drew et. al. [8] proposed a skull thinning technique in which glass is fused to the thinned skull providing a chronic and stable window with minimal bone regrowth. However, this window is not suitable for OR-PAM due to the acoustic impedance mismatch between glass and brain tissue [3]. Recently, a soft cranial window was employed for a ring transducer based OR-PAM brain imaging using agarose and acetate film to provide improved acoustic coupling [3]. However, the procedure for cranial window implantation required hours of anesthesia administration which can lead to brain alterations. Additionally, the utilization of ring transducer needed a large coupling medium which increased the complexity of the imaging. The bulky imaging head, also a characteristic of conventional OR-PAM setups, further limits its potential integration with traditional optical neuroimaging modalities [9–12]. Non-contact PAM techniques [13, 14] can potentially remove the need for coupling medium, but they require additional lasers therefore increasing cost and complexity of the setup. Recently, transparent ultrasound transducers (TUTs) based on transparent piezoelectric materials such as lithium niobate (LN) have been investigated for photoacoustic applications [15–17]. They provide a simple and cost effective solution as their transparency allows them to be placed co-axially in the optical path while being in close proximity of the biological target. This provides the capability to capture acoustic waves with minimal to no coupling medium requirement. However, engineering challenges remain to integrate TUT for OR-PAM brain imaging setups and demonstrate their potential for multimodal imaging.

To address the various issues mentioned above, we report a solution of an implanted LN-TUT window on thinned mouse skull for head restraint awake brain imaging with optical access. Our main idea follows Drew et. al. [8] thinned skull window process but replaces the glass with TUT. The implantation procedure required around sixty minutes and mouse recovered within a week after the procedure. This minimized any possible brain alterations. Moreover, the resultant simplified imaging head allowed for potential multimodal integration seamlessly. The implanted TUT window was observed to be stable over six weeks with minimal bone regrowth. Further, we employed the TUT for OR-PAM imaging. The TUT based OR-PAM setup was evaluated using a multilayered leaf phantom showing its depth sectioning capabilities. Next, we demonstrated in-vivo awake mouse brain vasculature OR-PAM imaging along with intrinsic optical signal imaging (IOSI) and twophoton microscopy (2PM) through the same TUT window indicating its potential for multimodal imaging.

The LN TUT was fabricated similar to previously reported literature [17]. In summary, indium tin oxide (ITO) coated 3 mm × 3 mm, 250 *µ*m thick LN (Precision Micro Optics, Burlington, MA, USA) was used as the piezoelectric material. This was followed by the backing layer composed of a transparent epoxy (EPO-TEK301, Epoxy Technologies Inc., Billerica, MA, USA). Two∼10 mm long gold pin terminated electrical wires were connected to the TUT using a silver epoxy (E-solder 3022,Von Roll Isola Inc., New Haven, CT, USA). Lastly, a 40 *µ*m thick Parylene-C layer was deposited on the transducer for acoustic matching and bio-compatibility.

For implantation, we followed a similar process outlined by Drew et. al. [8] but replaced the conventional cover micro glass slide by a 1 mm thick and 3 mm× 3 mm size 13 MHz LN TUT. C57BL/6J mice (male, 2–4 months old, Jackson Laboratory, Bar Harbor, ME, USA) were used to implant the TUT over the thinned skull for head restraint imaging. The surgical procedure was conducted as follows: a midline scalp incision was firstly made to expose the skull. Then a custom-made head bar was attached to the skull using cyanoacrylate glue (#32002, Vibra-tite, Clawson, MI, USA), and then sealed using dental cement (Ortho-Jet, Lang Dental, Wheeling, IL, USA). Next, in order to implant the TUT, a region of skull up to approximately 4 mm × 4 mm (Bregma to Bregma -4 mm and a width of ∼4 mm with sagittal sinus at the center) was thinned using a highspeed drill (K.1070 Micromotor Kit, Foredome, Bethel, CT, USA). Heat damage was prevented by constantly rewetting the skull surface with cold saline. Extra care was taken to not damage the sagittal sinus. The skull was thinned until to around tens of microns, where the pial vessels could be clearly observed. To implant the TUT, a thin layer of cyanoacrylate glue was placed on the thinned skull surface and TUT was then attached it. Lastly, all the wounds and edges were closed using the dental cement as schematically depicted in Figure 1(a). After the TUT window installation, the mouse was monitored for a week to allow for recovery and acclimation to the implanted head bar and TUT. Due to the natural curvature of the mouse skull, the implanted TUT needs to be adjusted to allow a normal illumination from the pulsed laser. Therefore, a custom built 6 degrees of freedom head restraint imaging setup, modified based on [19], was used to accommodate the TUT implantation angle as shown in Figure 1(b)-(c). The animal was fixed to the head restraint setup incrementally for 15, 30 and 60 minutes over three consecutive days to habituate to the setup before imaging sessions [20]. All surgeries and procedures in the experiment were carried out in accordance with animal use protocols approved by the Institutional Animal Care and Use Committees at the Pennsylvania State University as well as the guidelines established by the Use of Animals from the National Institutes of Health.

**Figure 1:**
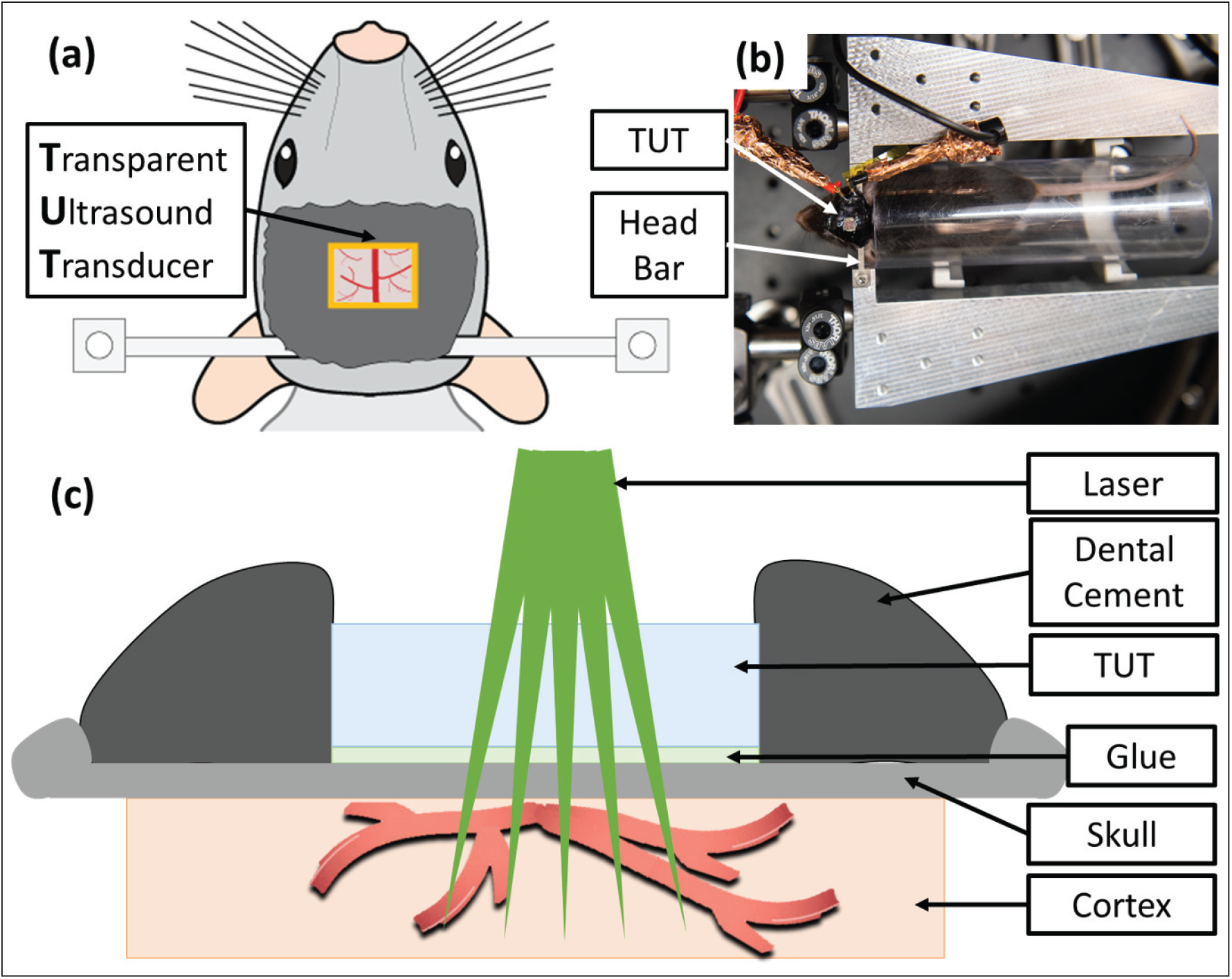
**(a)** Schematic of implanted transparent ultrasound transducer (TUT) window over mouse thinned skull and head bar placement. Image adapted from [18]. **(b)** The mouse head restraint imaging setup. The head restraint setup allows 6 degrees of freedom for alignment. **(c)** The sectional schematic of the mouse head TUT implantation.

The setup shown in Figure 2 was used for awake mouse brain OR-PAM imaging. A high-speed pulsed laser (GLPM-10, IPG Photonics; 532 nm wavelength; 1.4 ns pulse duration; 10-600 kHz repetition rate) driven at 10 kHz was employed and the output beam was sampled using a 10 % beam sampler (BSF10-A, Thorlabs Inc., Newton, NJ, USA) followed by photodiode (DET10A, Thorlabs Inc.). The detected beam samples were used to synchronize the system with the 16 bit, 1 Giga samples per second data acquisition system (Razormax-16, Dynamic Signals LLC, Lockport, IL, USA). The major beam passed through a neutral density filter (FW1AND, Thorlabs Inc., Newton, NJ, USA) and was redirected using mirrors to the spatial filter and collimating system consisting of a 20 *µ*m pinhole (P20D, Thorlabs Inc., Newton, NJ, USA) and 2 lenses, with focal lengths of 50 (LA1131-A, Thorlabs Inc., Newton, NJ, USA) and 75 mm (LA1608, Thorlabs Inc., Newton, NJ, USA) respectively. The collimated beam was focused using an achromat doublet lens (AC254-200-A, Thorlabs Inc., Newton, NJ, USA) with 200 mm focal length and steered using a two dimensional galvanometer scanner (GVS002, Thorlabs Inc., Newton, NJ, USA) for raster scanning the target through the TUT. The generated photoacoustic signals were filtered using a 3-30 MHz RF bandpass filter (ZABP-16+, Mini-Circuits, Branson, MO, USA) followed by amplification using two 28 dB amplifiers (ZFL-500LN+, Mini-Circuits, Branson, MO, USA) in series before being acquired by our data acquisition system. The steering mirrors in galvanometer were driven by a function generator (SDG1032X, SIGLENT Technologies, Ohio, USA) using ramp signals with 80% duty cycle and the laser was operated at 10 kHz. Additionally, data binning of 4 bit streams was performed to reduce background noise. The imaging speed i.e. B-scan and volumetric scan rate were 12.5 Hz and 0.03 Hz, respectively, for a 400 × 400 pixel image. The speed can further be increased by increasing the laser repetition rate.

**Figure 2:**
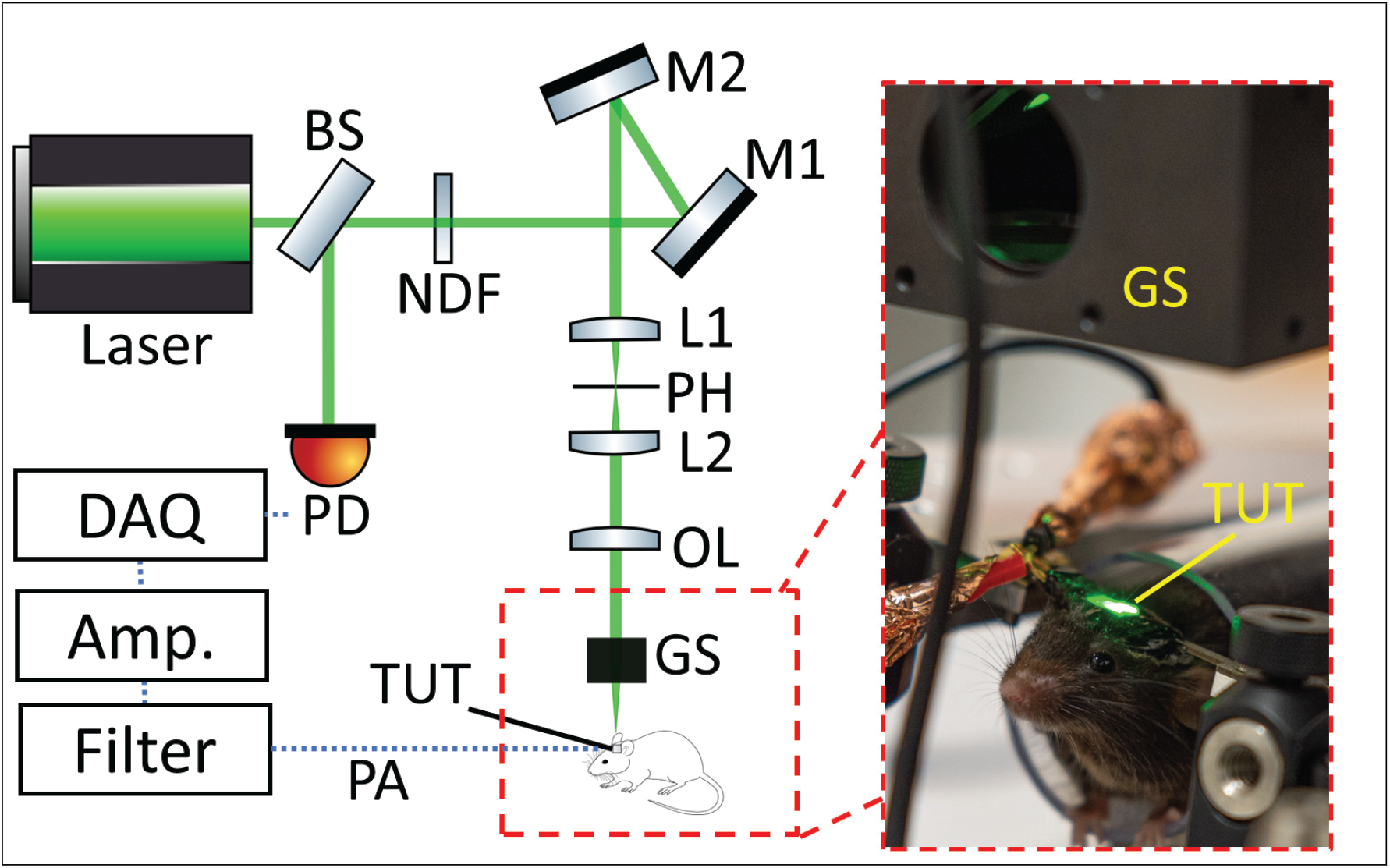
Experimental setup for awake mouse brain OR-PAM imaging using our TUT window. The photo shows the contents of the red dashed box, containing the imaging head with light being scanned using a galvanometer scanner without the requirement of an external coupling medium (see Visualization 1). BS: Beam splitter, NDF: Neutral density filter, M1: Mirror 1, M2: Mirror 2, L1: Lens 1, PH: Pinhole, L2: Lens 2, OL: Objective lens, GS: Galvanometer scanner, TUT: Transparent ultrasound transducer, PA: Photoacoustic signal, Amp: Amplifiers, DAQ: Data Acquisition system, and PD: Photodiode.

We first evaluated our TUT based OR-PAM setup’s resolution and imaging capabilities on phantoms. The lateral resolution was obtained by edge scanning a USAF resolution target shown in Figure 3(a). The edge spread function (ESF) was plotted along the red dashed line of the figure inset. The derivative of the ESF, which is the line spread function (LSF) is also plotted in the same figure. The lateral resolution was calculated from full width half maximum (FWHM) of the LSF, and was found to be 6.92 *µm*, indicating the setup’s ability to map microvasculature. Furthermore, by taking the FWHM of the Gaussian fitted photoacoustic axial line (Figure 3(b)), the axial resolution was calculated to be 202.5 *µm*.

**Figure 3:**
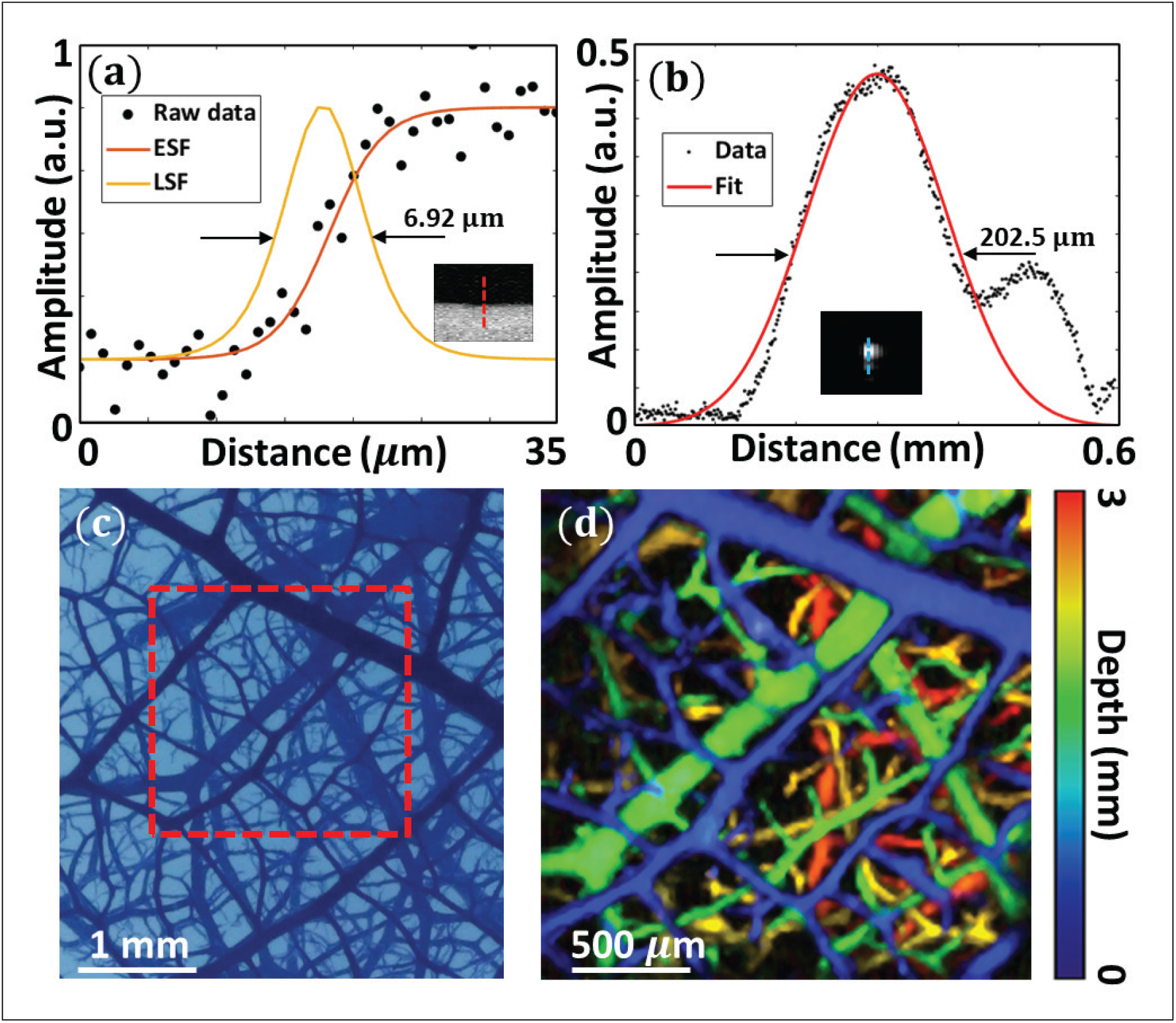
**(a)** Lateral resolution measurement from curve fitted edge spread function (ESF) and line spread function (LSF) obtained by scanning edge of USAF target (figure inset shows photoacoustic maximum intensity projection (MIP) image of the USAF target). **(b)** Gaussian fitted photoacoustic axial line of a 7 *µ*m carbon fiber (figure inset shows the B-scan of the carbon fiber). **(c)** Brightfield microscopy image of the multi-layered leaf phantom, the leaf has four layers within a span of 3 mm. **(d)** Reconstructed color-coded depth photoacoustic MIP image of the multilayered leaf phantom across an area of 2*mm* × 2*mm* highlighted by red dashed box in (c).

Next, we tested the imaging capabilities of our TUT based OR-PAM setup using a multilayered leaf phantom with four leaf layers spread across a 3 mm depth. The brightfield microscopy image of the phantom is shown in Figure 3(c). The leaf phantom was prepared in 1.5% agarose solution. The phantom was imaged over an area of 2 × 2 mm^2^ through the TUT window highlighted by the red box inset in Figure 3(c). The color-coded depth maximum intensity projection (MIP) photoacoustic image of the leaf phantom is shown in Figure 3(d). The reconstructed photoacoustic image faithfully followed the brightfield microscopy image and different layers were demarcated by different colors indicating different depths. Finer leaf structures up to 3 mm deep were able to be resolved, therefore demonstrating the setup’s overall ability to resolve fine structures while providing depth sectioning abilities.

Next, we head-fixed the mouse with TUT window implantation and imaged the vasculature of the awake mouse brain using the OR-PAM setup. The scanning was done through the TUT window and generated photoacoustic signal was captured by the TUT itself (Figure 2 and see Visualization 1). The MIP of the photoacoustic image is plotted in Figure 4(a) and the corresponding color-coded depth MIP photoacoustic image is shown in Figure 4(b). Photoacoustic responses from major vessels as well as their depth information were observed. The setup, limited by optical scattering, had a imaging depth of 0.4 mm. Furthermore, to demonstrate the multimodal imaging capability of our TUT window, we used two commonly employed neuro-optical imaging modalities: IOSI for wide field hemodynamics monitoring and 2PM for microcirculation monitoring. The setups and methods can be found in [18]. IOSI images were created by CCD camera capturing the vasculature reflectance from two collimated and filtered 530 ±

**Figure 4:**
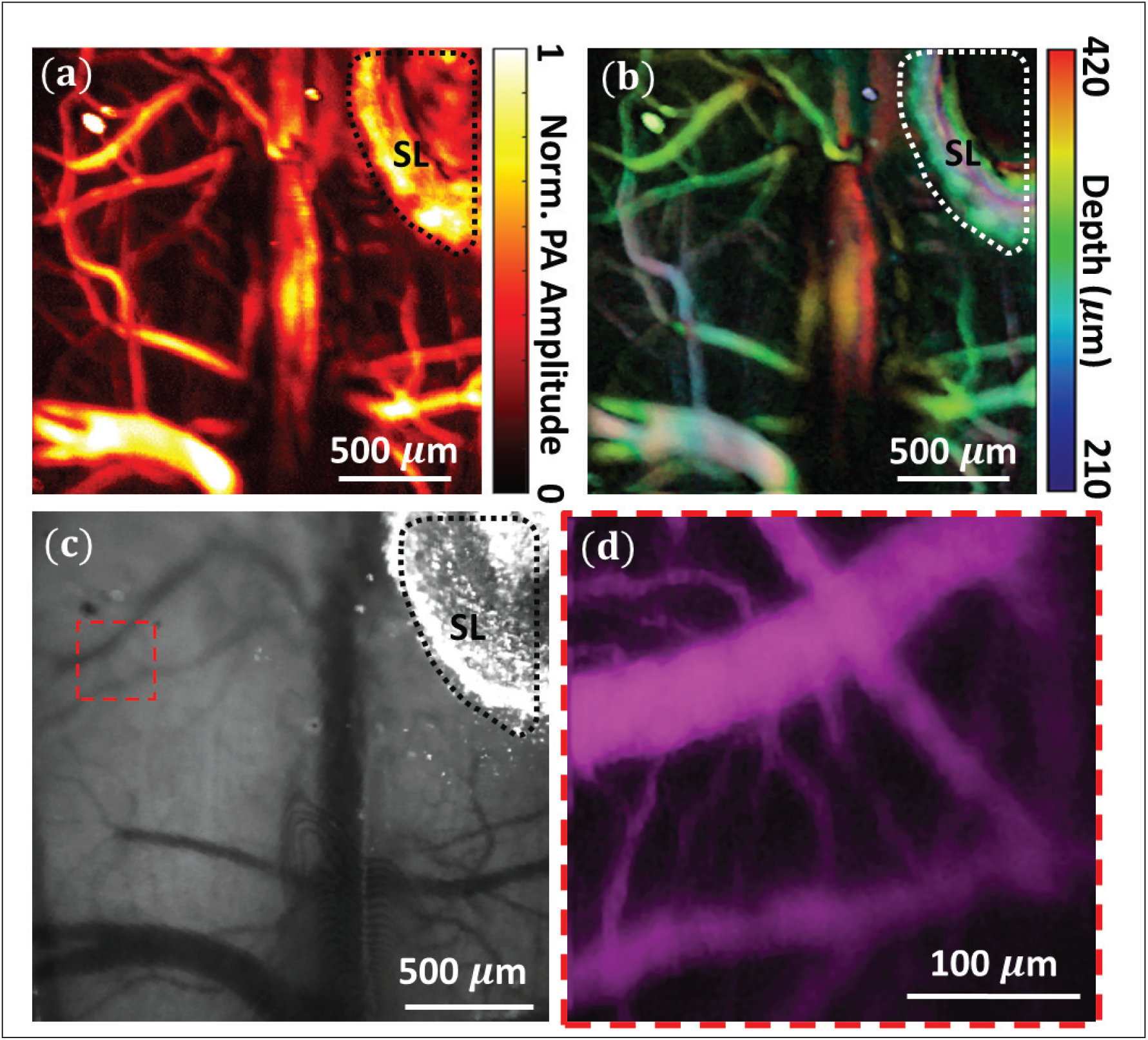
**(a)** Reconstructed maximum intensity projection (MIP) of the photoacoustic image in the x-y direction. **(b)** Color-coded depth MIP image of the photoacoustic image for the depth range of 0.21-0.42 mm from the transducer. **(c)** The mouse skull image through the implanted TUT window under LED illumination of the IOSI setup. The resolution of the captured image was 8.8 *µ*m/pixel. 2PM image of the red subset in (c) imaged through the TUT window, the field of view was 300 *µ*m × 300 *µ*m and the image was captured at a resolution of 0.825 *µ*m/pixel. SL: Silver Epoxy.

10 nm LEDs (FB530-10 filters Thorlabs Inc., Newton, NJ, USA, 10 nm FWHM). 2PM images were generated from injected fluorescein isothiocyanate–dextran (FITC) (FD150S, Sigma-Aldrich, St. Louis, MO) dyed vasculature, imaged with a 16X objective and an 800 nm Ti:Sapphire laser. Figures 4(c) and (d) shows the captured image of the IOSI and 2PM images through the implanted TUT window respectively. The captured IOSI image had a resolution of 8.8 *µm* per pixel and the major vessels and capillaries can be clearly seen through the TUT window. 2PM acquired fluorescence signal across 300 *µ*m × 300 *µ*m area, marked by red subset in Figure 4(c), with a resolution of 0.825 *µ*m per pixel. Finer capillary networks using 2PM imaging through the TUT window were clearly resolved. These results demonstrated the versatility of the implanted TUT window and its potential for awake mouse brain simultaneous multi-contrast imaging.

Besides serving as an ultrasound receiver and optical access window for multimodal imaging, in the future, the cranial TUT windows can also be used for delivering mechanical energy to the brain tissue for potential applications such as studying ultrasound induced neuromodulation and cancer drug delivery [21]. We recently demonstrated the capability of TUTs for high-throughput mechanical stimulation of cells while simultaneously imaging the induced calcium signaling with a high resolution confocal optical microscope [22]. The multi-functional aspects of our proposed cranial TUT window, therefore, could pave the way for a powerful awake mouse brain ultrasound stimulation platform combined with multimodal imaging capabilities.

## Supporting information

Laser scanning through the TUT cranial window for OR-PAM imaging in awake mouse

## Acknowledgments

We acknowledge Eugene Gerber and Mohamed Osman for support in the machining of parts and help with transducer design. We also thank Christopher Cheng for help in Parylene coating.

## Data availability

Data underlying the results presented in this paper are not publicly available at this time but may be obtained from the authors upon reasonable request.

